# Geographic microevolution of *Mycobacterium ulcerans* sustains Buruli ulcer extension, Australia

**DOI:** 10.1101/2020.11.03.366435

**Authors:** Jamal Saad, Nassim Hammoudi, Rita Zgheib, Hussein Anani, Michel Drancourt

## Abstract

The reason why severe cases of Buruli ulcers caused by *Mycobacterium ulcerans* are emerging in some South Australia counties has not been determined. In this study, we measured the diversity of *M. ulcerans* complex whole genome sequences (WGS) and reported a marker of this diversity. Using this marker as a probe, we compared WGS diversity in Buruli ulcer-epidemic South Australia counties versus non-epidemic Australian counties and further refined comparisons at the level of counties where severe Buruli ulcer cases have been reported. Analyzing 218 WGS (35 complete and 183 reconstructed WGS, including 174 Australian WGS) yielded 15 *M. ulcerans* complex genotypes, including three genotypes specific to Australia and one genotype specific to South Australia. A 1,068-bp PPE family protein gene exhibiting genotype-specific sequence variations was employed to further probe 13 minority clones hidden in sequence reads. The repartition of these clones significantly differed between South Australia and the rest of Australia. In addition, a significantly higher prevalence of 3/13 clones was observed in South Australia counties of the Mornington Peninsula, Melbourne and Bellarine Peninsula than in other South Australia counties. The data presented in this report suggest that the microevolution of three *M. ulcerans* complex clones drove the emergence of severe Buruli ulcer cases in some South Australia counties. Sequencing one specific *PPE* gene served to efficiently probe *M. ulcerans* complex clones. Further functional studies may balance the environmental adaptation and virulence of these clones.

## Introduction

Buruli ulcer, a handicapping skin and subcutaneous infection, was initially described in Australia along with the description of the first isolate of the causative *Mycobacterium ulcerans* (*M. ulcerans*).^1^ Thereafter, Buruli ulcers have been reported in several tropical areas of Asia, Africa and South America; therefore, 36 countries are currently notifying Buruli ulcer cases to the World Health Organization (WHO). In these endemic countries, the evolving incidence of Buruli ulcer is highly variable: the WHO is reporting a 64% decrease in notified Buruli ulcer cases worldwide over the past 9 years:^1^ in fact, the incidence of Buruli ulcer is decreasing in most West African countries, which declared the highest numbers of cases during the 1980-2000 period, whereas it is increasing in its birthplace, South Australia.^2,3^ The infection is specifically re-emerging in Victoria, southeastern Australia, extending to previously Buruli ulcer-free countries where severe forms are now encountered. ^2^ For West Africa, we previously reported a significant correlation between dropping incidence in Buruli ulcers and climate change, ^4^ while the reasons for the pejorative epidemiological evolution in Australia are unknown, although the hypothesis of currently undetected virulent clones of *M. ulcerans* has emerged. ^5,6^

Buruli ulcer is a non-transmissible disease contracted by direct or indirect contacts of breached skin with *M. ulcerans*-contaminated environments. ^1,7^ Therefore, the evolution of the incidence of Buruli ulcers in any one endemic area as well as its geographical extension in any one country, partly reflects the evolving dynamics of *M. ulcerans* in the ecosystems where this opportunistic pathogen is thriving. *M. ulcerans* is a close derivative of *Mycobacterium marinum* (*M. marinum*) after their common ancestor evolved following a strong reduction of the genome and the acquisition of a 174-210-kb giant plasmid, ^8^ encoding the nonribosomal synthesis of macrolide exotoxins named mycolactones. ^9^ *M. ulcerans* is repeatedly detected in Buruli ulcer-endemic tropical countries in stagnant water ecosystems to which rural populations that are developing Buruli ulcer are exposed;^10,11^ but only two environmental *M. ulcerans* isolates have been reported for which no genome sequence is available,^12,13^ meaning that the knowledge regarding the genomic diversity of the pathogen is limited to that of clinical isolates, which may not reflect the actual genomic diversity of *M. ulcerans* in the environment. Accordingly, the genomic microevolution of *M. ulcerans* clinical isolates has been reported in the Democratic Republic of Congo and Angola,^14^ but such observations were not placed in the background of evolving Buruli ulcers in the same geographic areas.

In this study, by analyzing the whole set of genomic data issued from studies of *M. ulcerans* in Australia,^15^ we reconstituted WGS data and observed the genomic microevolution of isolates specifically associated with the geographical expansion of *M. ulcerans* in coastal areas in South Australia^15^ and found one specific marker for that evolution.

## Materials and methods

### Isolate culture and sequencing

Total DNA of nine isolates (including 01 *M. marinum* B and 08 *M. ulcerans*) maintained in the Reference Mycobacteriology Laboratory, IHU Méditerranée Infection, France, was extracted from several colonies of a 6-week-old subculture on Coletsos culture medium using the InstaGene matrix following the manufacturer’s instructions (Bio-Rad, Marnes-la-Coquette, France). Then, the DNA (0.2 μg/μL) was sequenced using a MiSeq platform (Illumina, Inc., San Diego, CA, USA). DNA was fragmented and amplified by 12 cycles of PCR. After purification on AMPure XP beads (Beckman Coulter, Inc., Fullerton, CA, USA), the libraries were normalized and pooled for sequencing on a MiSeq instrument.^16^ Each sequencing run included one negative control (free of mycobacterial DNA) with ten mycobacterial samples to evaluate the quality of output sequence reads. Furthermore, data reads from different laboratories were used in this study.

### Genomic data analysis

In the first step, we retrieved WGS data for 44 *M. ulcerans* and *M. marinum* WGS (including the 09 WGS reported above), as well as inside reads deposited for 174 WGS made from *M. ulcerans* isolates in Australia (S1 Table).^2,3,15^ Then, the 182 *M. ulcerans* genome sequences and one *M. marinum* M were assembled and annotated using Spades version 3.14.0^17^ and Prokka version 1.14.6^18^, respectively. Roary 3.13.0^19^ was used with 95% identity to generate the pangenome, core genome and SNP-core genome of the 182 studied WGS (based on standard assembling software, such as SPAdes, A5-assembly and De novo). Moreover, PHYRE2 Protein Fold Recognition Server, CPHmodels-3.2, SCRATCH protein predictor (http://scratch.proteomics.ics.uci.edu/index.html) and Protter (http://wlab.ethz.ch/protter/start/) were used to predict the structure localization and function of genes of interest. CLC genomics 7 was used to map the output sequence reads against a gene and to detect the probability variant to estimate the zygosity (homozygous and heterozygous) of strains. Artemis software was used to visualize the heterogeneity profile of the studied strains (https://www.sanger.ac.uk/science/tools/artemis). WebLogo3 (http://weblogo.threeplusone.com/create.cgi) was also used to compare the gene sequences. We determined the properties of each genome (total length, G+C content, number of contigs and others) with Quast 5.0.2.^20^ Average nucleotide identity (ANI) was calculated using OrthoANIu^21^ to validate the classification results of *M. ulcerans* isolates. Moreover, Seaview version 4^22^ was used to view the alignment profile of the core genome and to detect differences in substitution mutations, deletions, or insertions between isolates.

## Results

### Whole genome analyses

We observed 41 different G+C% values among the 218 studied genomes, with G+C% values varying between a maximum of 65.8% for *M. marinum* strain DL240490 from Israel to a minimum of 63.73% for *M. ulcerans* SRR6346311 from Queensland, Australia and 63.37% for *M. ulcerans* CSURP9951 from Japan. Then, we observed that the genome size varied between a maximum length of 6.66 Mb for *M. marinum* strain M and a minimum length of 5.19 Mb for *M. ulcerans* SRR6346343 isolated from *Trichosurus vulpecula* in Gippsland, Australia (S1 Table). Studying ANI, core genome and SNPs based on core genome sequence (2,814-genes; 2,607,386-bp) data indicated six different *M. ulcerans* genotypes comprising 188 *M. ulcerans* isolates (A-F), one *Mycobacterium liflandii* genotype comprising two isolates (G), one *Mycobacterium pseudoshotii* genotype comprising four strains (H) and seven *M. marinum* genotypes comprising 24 isolates (I-O) (Figs 1a, 1b) (S2, S3 Tables). Focusing on the 174 *M. ulcerans* isolates made in Australia, we observed three different genotypes: genotype A included 163 clinical isolates, with SNPs varying between 1 and 950 (including isolates originating from six Australian counties, i.e., (Ballarat, Mornington Peninsula, Phillip Island, Melbourne, Gippsland and Bellarine Peninsula) and 06 animal isolates (01 *Canis lupus familiaris*, 03 *T. vulpecula*, 01 “koala”, 01 *Potorous longipes*) originated from four counties (Mornington Peninsula, Phillip Island, Gippsland and Bellarine Peninsula); genotype B included ten clinical isolates, with SNPs varying between 1 and 396 (including one isolate from Darwin, one isolate from Port Hedland and eight isolates from Queensland); and genotype C included one clinical isolate from Papua New Guinea grouped with *M. ulcerans* CSURQ0185 isolated from Côte d’Ivoire with 383 SNPs (S2 Table). We found that the values of SNPs between the genotypes were as follows: between genotypes A and B >1,000, between genotypes B and C >2,000 and between genotypes A and C >2,000.

**Fig 1:**
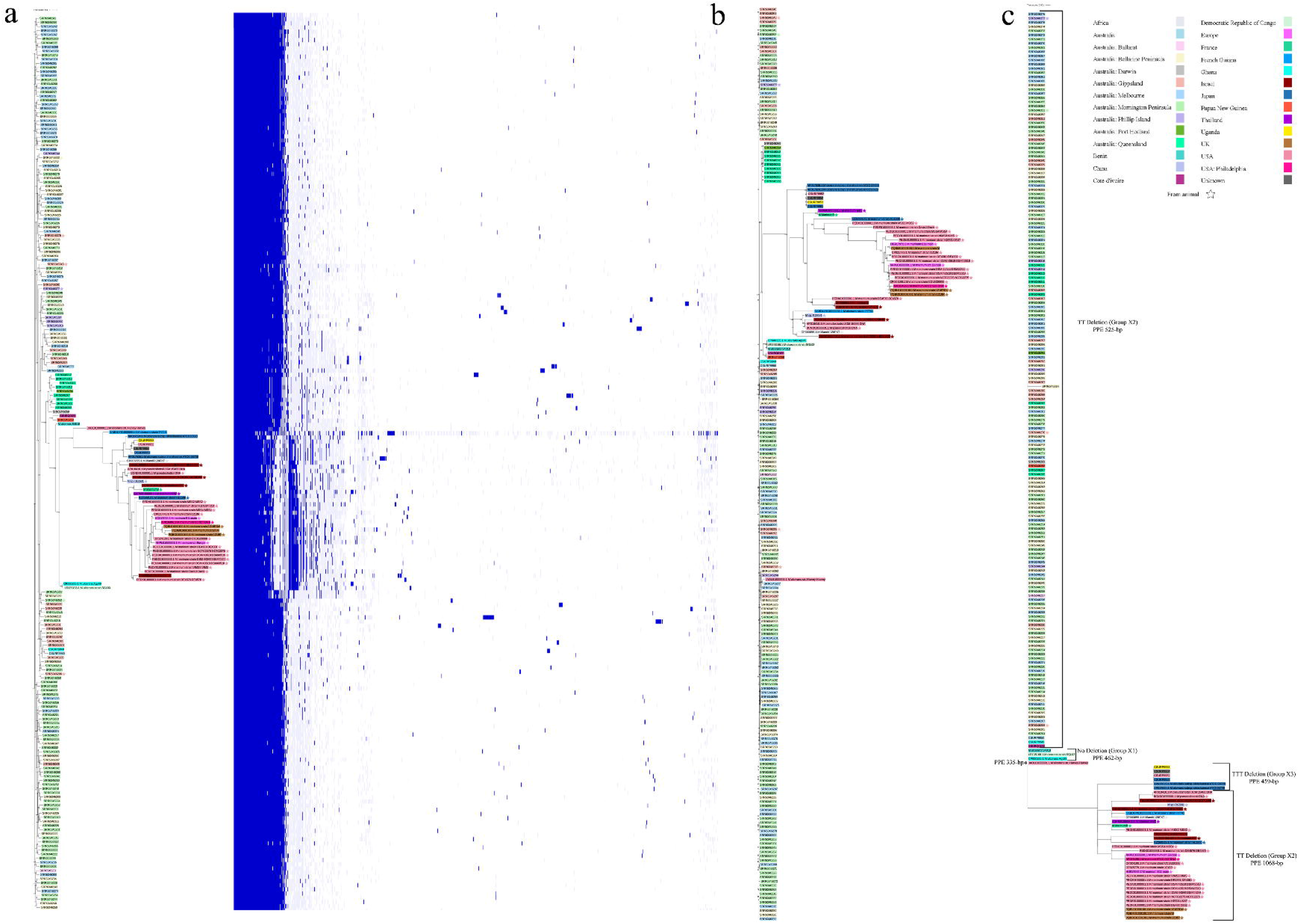
a: Pangenome tree with profile genes of 218 studied strains; b: core-genome tree of 218 studied strains; c: *PPE* gene tree of 217 strains with the gene length for each isolate. The three groups X1, X2 and X3 of the studied strains are indicated. Different colors were used to show the isolation counties, and a star was used to mark animal isolates from another.

### Hotspot-specific target

Core genome sequence alignment of the 218 genomes examined in this study (Fig 1b) indicated that one 1,068-bp *PPE* gene was annotated as a PPE family protein in the *M. ulcerans* strain CSURP7741 reference genome (GenBank assembly accession number GCA_901411635.1). The PPE protein was predicted to be an alpha helical transmembrane protein with 0.934205 probability. This gene exhibited interesting heterogeneities, including that it was completely absent in *M. ulcerans* SRR6346343 isolated from *T. vulpecula* in Gippsland. First, we observed deletions in the 5’ extremity of the gene resulting in a *PPE* gene size varying between 335 bp and 1,068 bp, correlating with the clusterization of the isolates in the core-genome tree (Fig 1b and 1c). These PPE deletions resulted in a non-transmembrane protein prediction with 0.90177 probability. Second, we observed a heterogeneous sequence hotspot sorting 216 isolates into three groups (this hotspot region was absent in *M. ulcerans* strain Harvey (JAOL01000001.1) and in *M. ulcerans* (SRR6346343). Group X1 was characterized by no deletion variant *PPE* gene at 376-377-378-379-380-381 (AATTTT) sequence position in three *M. ulcerans* isolates; group X2 was characterized by a TT deletion at positions 378-379 in 177 *M. ulcerans* isolates, in all four *M. pseudoshotii* isolates, in the two *M. liflandii* isolates and in all 24 *M. marinum* genotypes; and group X3 was characterized by a TTT deletion at positions 378-379-380 in six *M. ulcerans* isolates (Fig 2, S1 Table). Focusing on the 173 *M. ulcerans* isolates from Australia, we found that all these isolates had a 525-bp group X2 *PPE* gene (Fig 1c).

**Fig 2:**
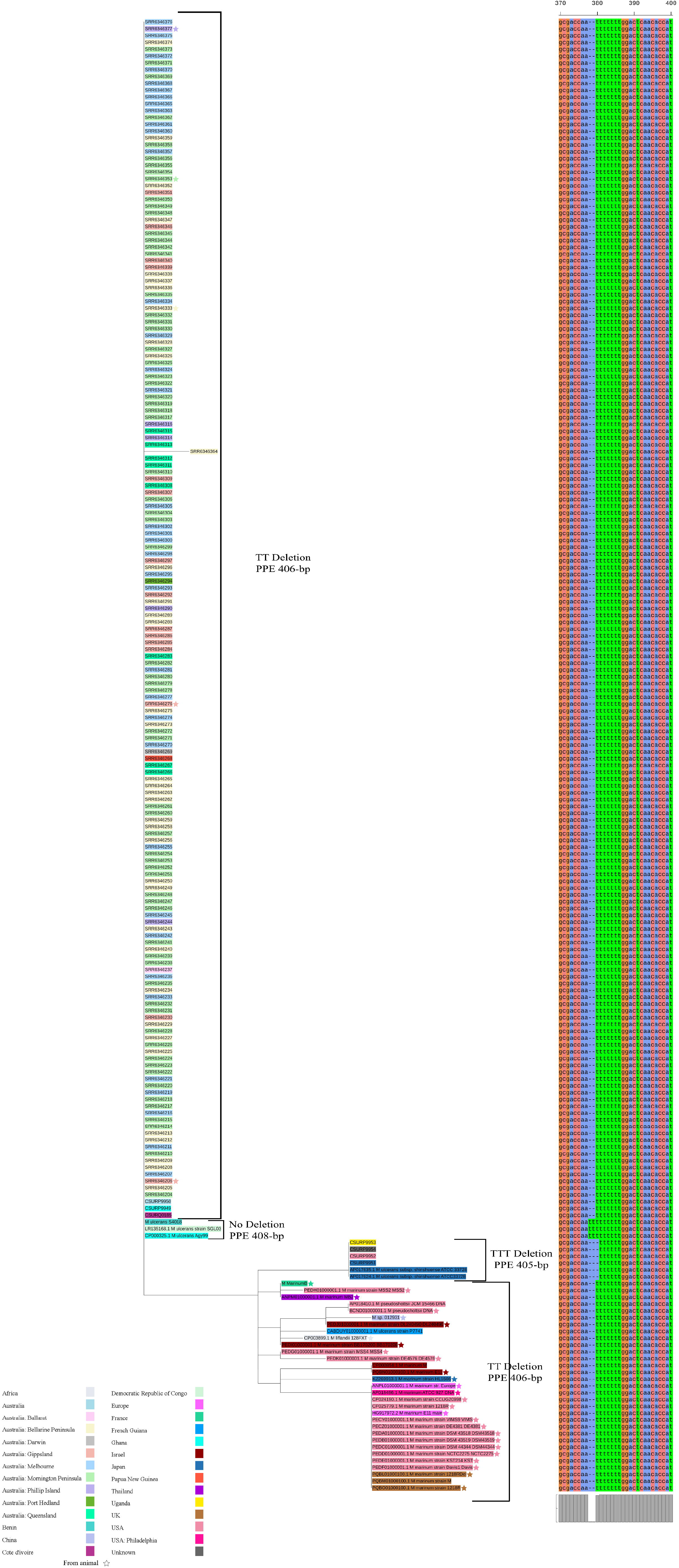
Tree based on the common region of the *PPE* gene of 216 studied strains with the different groups X1, X2 and X3 depending on the *PPE* gene variant (at positions 378, 379 and 380).

### *M. ulcerans* diversity, Australia

To question the clonal diversity of the Australian isolates of *M. ulcerans*, we mapped the output MiSeq reads of 182 genomes (issued from 174 Australian isolates plus eight isolates sequenced in IHU Méditerranée Infection, France) to the hotspot region. We searched for the specific target deletion in the depth of reads and found additional deletion forms. In total, 13 *PPE* variants (here designed PPE-1, PPE-13) were detected among the 182 isolate genomes, including PPE-1 [AATTTT], PPE-2 [AA--TT], PPE-3 [AA-AAT], PPE-4 [AA-ATT], PPE-5 [A--CTT], PPE-6 [AA-TTT], PPE-7 [AAATTT], PPE-8 [AA---T], PPE-9 [--TATT], PPE-10 [AA--CT], PPE-11 [AA--AT], PPE-12 [AA--GT] and PPE-13 [AA-CTT], in addition to the three variants reported above (Fig 3). As an example, we extracted corresponding reads of *M. ulcerans* strain CSURP9950, mapped them against hotspots using Bowtie software,^23^ and then we performed a reassembly. As a result, we obtained two different sequences: a majority sequence (TT deletion) and a minority sequence characterized by one T deletion followed by one A substitution; both sequences exhibited only 92% sequence identity with one another. BLASTn results of these two sequences showed that the majority sequence matched with *M. ulcerans* group X2 with 100% identity, while the minority sequence matched with *M. ulcerans* group X1 with 99.2% identity. Furthermore, we observed that the *M. ulcerans* strain CSURP7741 lacked heterogeneity in the hotspot (S1 Fig). Overall, deep analysis yielded ten additional variants in the *M. ulcerans PPE* gene, updating to 13 variants in this gene (Including the 03 variants found by assembly genome analysis plus the 10 variants found by deep sequence analysis). Focusing on the 174 *M. ulcerans* isolates from Australia, we observed geographical clusterization within 11 clusters (S2 Fig), and in Victoria, we had 695 variants for 157 clinical isolates as follows: in Mornington Peninsula, we had 305 variants for 66 clinical isolates; in Melbourne, we had 177 variants for 37 clinical isolates; in Bellarine Peninsula, figures were 149/36; in Phillip Island, 14/4; in Gippsland, 41/13; and in Ballarat 9/1. Outside Victoria, we had 32 variants for 11 isolates as follows: Queensland, 23/8; Papua New Guinea, 2/1; Port Hedland, 4/1 and Darwin, 3/1 (S4, S5 Tables). Analyzing these output data by comparing the average of detected variants between isolates belonging to each counties using the Wilcoxon test indicated a significantly (p-value = 0.001834) higher polyclonal diversity of *M. ulcerans* isolates from Victoria, Southeast Australia than from North Australia (S6 Table). Moreover, inside Victoria, the analysis indicated that *M. ulcerans* isolates collected from the Bellarine Peninsula, Melbourne and Mornington Peninsula exhibited a significantly higher polyclonal diversity than *M. ulceran*s isolates collected from the other two Victoria counties (Gippsland and Phillip Island), with p-value = 0.02062, p-value = 0.0006314 and p-value = 0.0009121, respectively (S6 Table). Finally, we observed a significantly higher incidence of [AA-TTT], [AAATTT] and [AA--AT] variants in the three Buruli ulcer epidemic counties, Bellarine Peninsula, Melbourne and Mornington Peninsula^3,15^ compared to the non-epidemic counties using the chi-square (χ2) test with p-value=0.001367, p-value=0.0002324 and p-value=0.04721, respectively (S7 Table).

**Fig 3:**
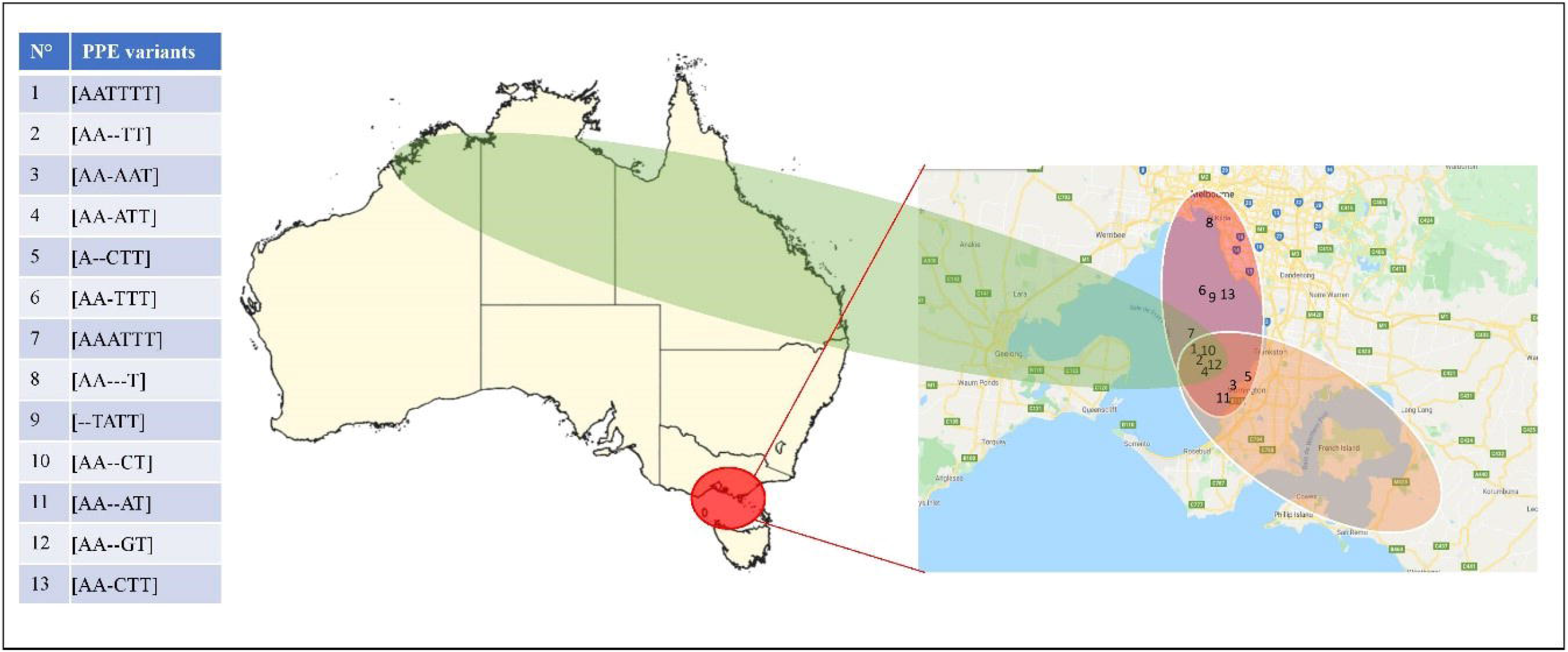
Map showing the distribution of the different *PPE* variants in Australia. The table shows the 13 *PPE* variants (see Text), the Venn diagram shows the *PPE* variants found in the endemic area in Victoria county (red), the non-endemic area in Victoria county (orange) and the non-endemic area outside Victoria county (green).

For animal isolates in Australia, we found 04 variants in 03 *T. vulpecula* isolates made from the epidemic counties Mornington Peninsula and the non-epidemic counties Gippsland and Phillip Island. Moreover, we found 03 variants in one *C. lupus familiaris* isolate made in one epidemic counties of the Bellarine Peninsula, 02 variants from one *P. longipes* isolate made in the non-epidemic counties of Gippsland and 03 variants in one “koala” isolate obtained from the non-epidemic counties of Gippsland (S2 Fig, S4 Table). We observed that the number of detected *PPE* gene variants in animal isolates was lower than that of human isolates from the same counties. Specifically, 4/13 *PPE* variants were detected in common in these four animal species (with 03 of them being endemic to Australia), whereas 9/13 *PPE* variants were found exclusively in *M. ulcerans* isolates of human origin (S4 Table). In summary, [AA--TT], [AA-ATT], [AA--CT] and [AA--GT] variants were found in common in animals and humans in both Buruli ulcer epidemic and non-epidemic counties in Australia (S4 Table).

## Discussion

In this study, reviewing *M. ulcerans* WGS data indicated that we and colleagues routinely deposited in public genome sequence databases only one straight genome sequence per *M. ulcerans* isolate, although such WGS data were derived from cultured colonies (i.e., millions of cultured mycobacteria), as single-cell sequencing has never been used in that context to the best of our knowledge.^24^ To overcome this limitation, we analyzed the entire set of available sequencing reads, instead of allowing any software to automatically analyze majority reads and ignore minority reads.^24,25^

While the resulting sequence heterogeneity may have resulted from contamination and sequencing bias, it may also carry biologically relevant information, as illustrated in this report.^26,27^ Following this original pathway of WGS analysis, we indeed observed a significantly higher genomic diversity among *M. ulcerans* isolates collected in Buruli ulcer endemic counties of South Australia compared to non-endemic ones. We were even able to refine the analysis to the epidemiologically relevant scale of Australian counties of the Mornington Peninsula, Melbourne, and Bellarine Peninsula, where a pejorative evolution of the infection has been recently reported.^2,3,15^ At this scale, we noticed that *M. ulcerans* clones tagged by specific *PPE* gene variants were detected in clinical samples in common with animal samples, three of which were found only in Australia.

The genomic data reported in this study indicate a roadmap for further studies aimed at translating genomic characteristics into microbiological characteristics of *M. ulcerans* isolates in Australia to precisely support the increased incidence and virulence of Buruli ulcer in some counties of southeastern Australia.

We propose generalizing the genome analysis described in this report to highlight minority clones generally obscured by majority clones in routine analyses, thereby helping to depict the biological diversity of medically relevant microorganisms, including opportunistic pathogens of environmental origin, such as *M. ulcerans*.

## Acknowledgments

We would like to thank the Department of Microbiology and Immunology, Doherty Institute for Infection and Immunity, University of Melbourne, Melbourne, Victoria, Australia and Doherty

Applied Microbial Genomics, Department of Microbiology and Immunology, Doherty Institute for Infection and Immunity, University of Melbourne, Melbourne, Victoria, Australia, for agreeing to publish the assembly genome of 174 clinical *M. ulcerans* isolates. J.S., N.H., R.Z. and H.A. are supported by a Ph.D. grants from the IHU Méditerranée Infection, Marseille, France.

## Authors’ addresses

Jamal Saad, Nassim Hammoudi, Rita Zgheib, Hussein Anani, Michel Drancourt. Aix-Marseille-Univ., IRD, MEPHI, IHU Méditerranée Infection, Marseille, France.

## Supporting information

**S1 Fig**. Heterogeneity profile at the level of depth reads of several isolates based on 125 bp of the PPE gene, including sequence study variants.

**S2 Fig**. Heatmap and hierarchical clustering tree showing the profile of 174 Australian isolates based on the presence or absence of the 13 detected variants and the isolation region of isolates.

**S1 Table**. Genomic and clinical information’s about isolates study.

**S2 Table**. Pairwise SNP distance matrix values between 218 studied strains.

**S3 Table**. Pairwise ANI values between 218 studied trains

**S4 Table**. Distribution of the different *PPE* variants in the different Australia counties.

**S5 Table**. Distribution of the different *PPE* variants in each isolates study of Australia.

**S6 Table**. Statistical chi-square (χ2) test of detected variants between Victoria counties and outside Victoria in Australia.

**S7 Table**. Statistical chi-square (χ2) test of variants between Victoria counties and outside Victoria in Australia.

